# Dynamic cognitive differences between internal and external attention are associated with depressive and anxiety disorders

**DOI:** 10.1101/2024.09.26.615274

**Authors:** Taiki Oka, Yutaro Koyama, Akihiro Sasaki, Misa Murakami, Nao Kobayashi, Aurelio Cortese

## Abstract

The internal/external attention framework characterises attention focused on internal representations, such as emotions, versus external representations, such as perceptual stimuli. The inability to focus one’s attention is considered a critical factor in psychiatric disorders. While these different attentional foci are likely generated by the dynamic interplay of multiple cognitive processes, previous studies have generally examined single cognitive dimensions. We developed a new method, cognitive dynamic similarity analysis (C-DSA), to clarify how cognitive processes differ between experimental conditions. In an MR scanner, participants performed a word-processing task in which they focused on either their own emotions or the number of letters associated with a stimulus. To extract cognitive dynamics at the single-trial level, we applied cognitive dynamics estimation, a recently developed method that generates whole-brain activation maps for four cognitive dimensions (emotion processing, selective attention, self-referential thought, and working memory) using a meta-analytic platform. We then performed C-DSA to calculate the difference between internal and external attention for each cognitive dimension. C-DSA revealed significant differences between internal/external attention in all cognitive dimensions, but especially in emotion processing. Moreover, the difference between attention conditions of selective attention was negatively associated with the severity of depression and state-anxiety, but positively associated with trait-anxiety. Our findings suggest that C-DSA applies to both naturalistic and controlled dynamic processes and may be valuable in clinical settings by linking dynamic cognitive mechanisms with issues like ageing and psychiatric disorders.

**Graphical abstract:** 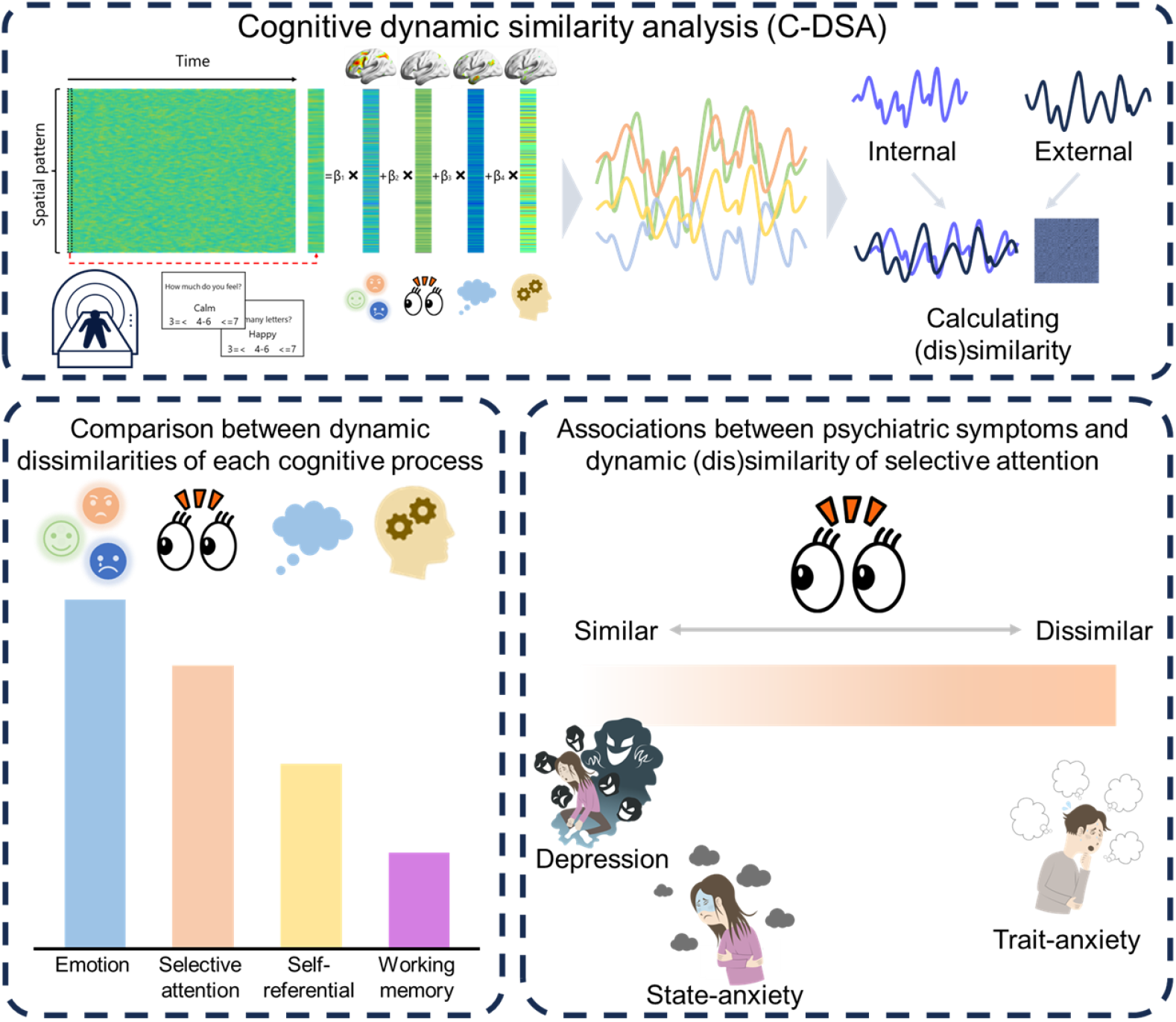

## Introduction

The internal/external attention framework characterises attention focused on internal representations, such as emotions, vs perceptual stimuli (Chun *et al*., 2011). Dysfunctions related to internal/external attention are thought to be present in many psychiatric disorders (Rafi *et al*., 2023). Previous research suggests that people with severe psychiatric symptoms, such as rumination, might have more difficulty than healthy people in dividing their attention between internal and external attention (Apazoglou *et al*., 2019). In response to these difficulties psychiatric patients face, intervention methods such as the attention training technique (ATT) have been proposed (Fergus & Wheless, 2018; Kowalski *et al*., 2020). However, internal and external attention are constructed from a complex and dynamic interplay of several cognitive processes, e.g., emotions, working memory, selective attention, etc (Mrkva *et al*., 2020; Lim & Pratt, 2023). Although many have stressed the importance of clarifying dynamics of various cognitive dimensions in internal/external attention and their alterations in psychiatric disorders, this area remains understudied (Amir & Bernstein, 2022; Morris & Braver, 2022; Verschooren & Egner, 2023). Understanding these dynamics is necessary to model maladaptive mental dysfunctions and to provide evidence for existing psychotherapeutic interventions.

To explore which cognitive processes are involved in specific functions and to investigate the nature of their involvement, fMRI has been used to investigate associations between brain regions and their activities. This is conventionally accomplished with regression analysis of brain activity. Representational similarity analysis has also been applied to compare brain representations between two conditions related to cognitive function (Freund *et al*., 2021). However, estimated brain maps based on this traditional approach cannot clarify individual processes when exploring temporal dynamics of complex functions, such as internal/external attention, constructed from various, overlapping cognitive processes. Furthermore, it is difficult to examine temporal dynamics of cognitive processes and their similarities in each type of attention. Developing a novel method is therefore essential to explore how dynamics differ between attentional types.

Here, we attempted to clarify the cognitive dynamic representation using an introspection task and a novel analytical method called cognitive dynamic similarity analysis (C-DSA, see Fig. 1). In C-DSA we compute the dissimilarity across cognitive processes, which is an index reflecting the difference between two complex cognitive dimensions. C-DSA allows us to investigate how independent cognitive processes differ between the two types of attention. It also enables us to investigate the relationship between these cognitive patterns and psychiatric disorders. Theoretically, psychiatric symptoms are related to an inability to divide/switch internal and external attention (Amir & Bernstein, 2022). Therefore, we hypothesised that the similarity of dynamics between internal/external attention should correlate with the severity of each illness.

**Figure 1.**
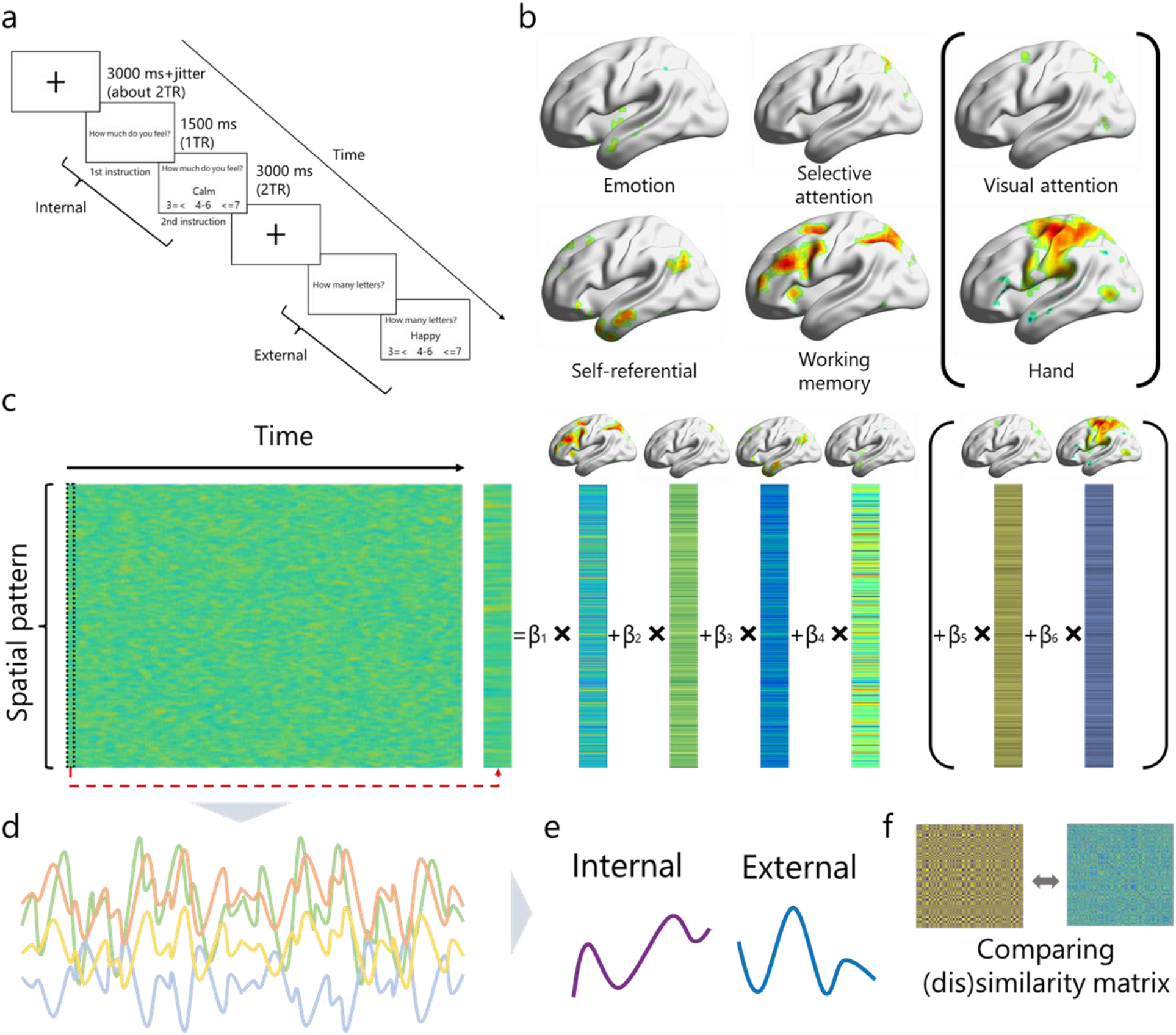
Scheme of cognitive dynamic similarity analysis for an internal/external introspection task. (a) The procedure for the introspection task. (b) Brain activation mapping extracted from meta-analytic software (Neurosynth (Yarkoni et al., 2011)) (c) The image of transposed GLM based on cognitive dynamical estimation (CDE) (Koyama et al., 2023) (d) Obtaining time series derived from coefficients of each cognitive process (e) Dynamic dissimilarity matrices (DDM) and making comparisons. (f) Creating the DDM and calculating the distance correspondence index (DCI) by comparing the distance matrix of attentional types and DDMs of each cognitive process. See Methods for details of the C-DSA.

If such cognitive dynamic processes can be clarified, they may tell us about cognitive pathways underlying difficulties with critical competencies related to self, including both internal attention and emotional dysregulation in psychiatric disorders. Results of the present study indicate that dynamics of an *a priori* cognitive process, which were chosen based on previous evidence for internal/external attention, differed between attentional types. We also found that these similarities are associated with psychiatric disorders, especially depression and anxiety. Our findings clarify how dynamics of each cognitive function are linked to psychiatric symptoms, with clinical implications for psychological interventions and neurobehavioural therapeutics.

## Materials and Methods

### Transparency, openness, and ethics

The current study has not been pre-registered. Data, relevant scripts for all analyses, and other supplementary materials may be found at open science framework (OSF, DOI:10.17605/OSF.IO/CQKNZ). The supplementary methods also contain detailed task instructions and stimuli employed. This study was approved by the Ethics Committee of the Advanced Telecommunications Research Institute International (Japan) (No.182H).

### Participants

We recruited 19 participants (mean age 22.8 y.o.; SD ± 1.64; 3 females) from the general population. One participant was excluded because the number of correct answers in the external trial was around chance level (33.8%, *p* = 0.86 binomial test against chance level of 33%). As a result, 18 participants (mean age 22.8 y.o.; SD ± 1.64; 2 females) were analysed. All had normal or corrected vision using MRI-compatible glasses. All participants provided written consent before beginning the experiments.

### Introspection task

We conducted an introspection task that required participants to focus on internal/external attention during a word-processing task regarding emotion (Fig 1a, Apazoglou *et al*., 2019; Rafi *et al*., 2023). Each trial began with a 1.5-s instructional screen indicating whether the subsequently presented stimulus required internal or external attention, followed by 3 s of stimulus presentation. At the bottom of each stimulus presentation screen, a numerical scale was displayed to help participants respond to the task. In internal trials, participants were asked to indicate the extent to which they were currently feeling the state indicated by an emotional word, e.g., calm, on a numerical scale of ≤ 3 (not feeling the indicated state or feeling it a little), 4-6 (feeling the indicated state moderately), ≥ 7 (strongly perceive the indicated state). In external trials, participants had to indicate the number of letters for a word, e.g., in the case of “calm,” 4-6 is correct. This response design allowed us to have the same design and stimulus presentation for both internal and external attentional trials. After each trial, a fixation cross with a jittered duration between 3,500 and 4,500 milliseconds was shown. Emotional words were extracted from the emotion and arousal checklist (EACL, (Oda *et al*., 2015) rather than from measures used in previous studies that employed a similar task (Apazoglou *et al*., 2019; Rafi *et al*., 2023), because their Japanese version was not appropriate for the length of words used as instructional material. The list of stimuli and examples of actual Japanese stimuli are shown in Table S1 and Fig. S1. Each task block lasted about 6 min, and participants completed eight blocks. Importantly, in previous studies using similar tasks (Apazoglou *et al*., 2019; Rafi *et al*., 2023), reaction time was controlled by instructing participants to “respond as quickly as possible,” but that was not done in this study. The reason for this more lenient instruction related to reaction time was to improve ecological validity and extract more natural dynamics.

### Clinical assessment of psychiatric disorders

We assessed the severity of multiple psychiatric scores using validated questionnaires. Each participant was evaluated for major depressive disorder (BDI, (Dozois *et al*., 1998)), obsessive-compulsive disorder (OCI, (Foa *et al*., 1998), internet-related problems (CIUS, (Meerkerk *et al*., 2009), attention-deficit/hyperactivity disorder (ADHD) (ASRS, (Kessler *et al*., 2005), autistic spectrum disorder (AQ, (Baron-Cohen *et al*., 2001), social anxiety (LSAS-fear/avoid, (Baker *et al*., 2002), state-trait anxiety (STAI-Y1&Y2, (Spielberger, 1983), and internet gaming disorder (IGDS-SF, (Lemmens *et al*., 2015). We also used the behavioural inhibition/activation system (BIS/BAS) questionnaire, but did not include it in this analysis because it was collected for a different purpose. Details of all questionnaires can be found in the Supplementary Methods. Importantly, we collected multiple psychiatric questionnaires because existing research often focuses on one or two psychiatric dimensions at a time, though the issue of internal/external attention is considered to span multiple psychiatric disorders.

### fMRI acquisition

The Psychophysics Toolbox for Matlab (http://psychtoolbox.org/) was used to conduct the experiments. A 3.0 T scanner (Prisma; Siemens, Erlangen, Germany) with a 64-channel head coil was used to collect fMRI neuroimaging data. We scanned 76 interleaved, axial, contiguous 2-mm slices, parallel to the anterior and posterior commissure line, using a T2*-weighted gradient-echo multiband echo-planar imaging (MB-EPI) sequence [repetition time (TR) = 1500 ms, echo time (TE) = 30.0 ms, flip angle (FA) = 70°, field of view (FOV) = 200 × 200 mm^2^, resolution = 2 × 2 mm^2^, MB factor = 6, voxel size = 2 × 2 × 2 mm^3^]. We obtained 240 volumes for the task. We took additional dummy images at the beginning of scanning. All individuals underwent a magnetization-prepared rapid acquisition gradient echo technique to acquire high-resolution T1-weighted images of the entire brain for anatomical reference. (MPRAGE, TR = 2250 ms, TE = 3.06 ms, FA = 9°, FOV = 256 × 256 mm^2^, voxel size = 1 × 1 × 1 mm^3^). Participants operated a response pad attached inside the MRI scanner with an adhesive sheet positioned to their right sides.

### Data analysis

#### Behavioural preprocessing

Trials with reaction times more than 3 SDs the mean, and/or that did not respond within the time limit, and/or that had incorrect responses in external attention trials, were excluded from the analysis. Trials with RTs less than 200 ms were also excluded.

#### fMRI preprocessing

Blood oxygen level dependence (BOLD) signals in native space were pre-processed in MATLAB (R2020b) (MathWorks) using SPM12 and in-house code. Images underwent motion correction, reorientation, and realignment with the first volume. Subsequently, T1-co-registered volumes were normalised using an MNI template. A linear trend was removed from time courses. Moreover, grand-mean intensity normalisation was performed to minimise baseline differences across blocks. No spatial or temporal smoothing was applied.

#### Cognitive dynamical estimation (CDE)

To reveal dynamics of each cognitive function in internal/external trials, we used cognitive dynamics estimation (CDE) (Koyama *et al*., 2023). CDE is based on linear regression of a two-dimensional fMRI matrix with spatial patterns of cognitive processes obtained from a meta-analysis as regression variables (Fig 1c). CDE can extract a time series of cognitive processes reflecting trial-wise cognitive processes. Using meta-analytic software, Neurosynth (Yarkoni *et al*., 2011), we generated activation maps of*a priori* four cognitive processes (Fig 1b):*emotions (Emo), selective attention (SA), self-referential thoughts (SR)*, and*working memory (WM)*. These processes were chosen because the cognitive processes are considered important for introspection, switching between external and internal representation (Apazoglou *et al*., 2019; Amir & Bernstein, 2022; Rafi *et al*., 2023; Gresch *et al*., 2024). Moreover,*visual attention (VA)* and*hand* processes were also chosen as no-interest regressors that might act as confounders. We confirmed the variance inflation factor (VIF) to assess multicollinearity, and there were no independent variables whose VIF was greater than 2.5 (Johnston *et al*., 2018). The model equation was defined as follows:

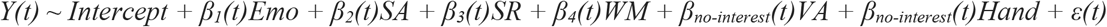

*Y*(*t*) represents the spatial pattern of fMRI data at time-point (*t*). As a result, we obtained a time series for each cognitive process during the task (the averaged time series through the sessions is shown in Fig S2). Then, we extracted each internal/external duration (5 volumes) centred on the second instruction from each trial after shifting by 6.00 s (4 TRs) to account for the hemodynamic delay. To confirm the validity of the analysis, we tested whether the local maxima of*hand* dynamics captured response timing using a binomial test and the performance was significantly higher than the chance level (average 32.1%,*p* < 0.001).

#### Cognitive dynamic similarity analysis (C-DSA)

Applying CDE, we obtained trial-wise time-series matrices of each cognitive process, excluding no-interest confounders as above. To compute representational maps of attentional direction (internal/external), we calculated the trial-wise dissimilarity (1-Pearson’s r) between internal/external attention. Then, for statistical analysis of differentiation in internal/external attention, we computed a distance correspondence index (DCI) to measure the relationship between each cognitive dynamic dissimilarity and attentional direction (Chikazoe *et al*., 2014). DCI was computed using a generalised linear logistic regression to predict the dissimilarity of attentional direction (0 or 1), which included all four independent variables, corresponding to dynamical distance in*emotion, selective attention, self-referential thoughts, working memory*, and regressors of no interest (differences in*visual attention* and*hand*). The formula to extract DCI is as follows:

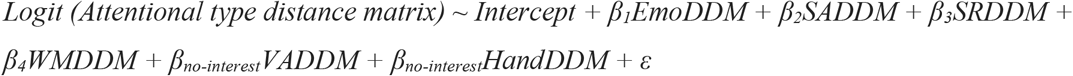

*DDMs* mean dynamic dissimilarity matrix. The extracted DCI as regression coefficients (*β*_*m*_) represents the extent to which the distance in each of the four cognitive properties predicted the dynamic similarity of attentional direction. DCIs for each cognitive process were computed for each participant and then submitted to the following statistical analysis.

#### Statistical analysis

For behavioural analysis, we tested RT differences between internal and external conditions using mixed-effect generalised linear regression after logarithmic transformation of RT data to verify differences in conditions. We also tested whether there were any differences in RT between repeated and switched trials because there were several consecutive trials of internal/external attention and switches between the types through each session. Participant effects were included as random effects in all tests.

Next, we conducted Wilcoxon signed-rank sum tests to examine whether each DCI was significantly above zero, which precisely revealed the difference in internal/external attention. In addition, we used the same test to compare the difference between cognitive properties, e.g., DCI of*emotion* vs. DCI of*selective attention*.

Moreover, we used mixed-effect generalised linear regression to reveal how each cognitive dynamic dissimilarity is related to psychiatric disorders. Each psychiatric score was input as an independent variable, and each cognitive time-series dissimilarity was input as a dependent variable, as follows:

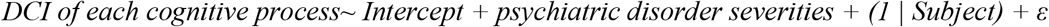

Finally, we obtained four model results for each cognitive process. Each regression coefficient represents how each psychiatric severity was associated with the degree of differentiation between internal and external attention in each cognitive process.

For all statistical analyses, a*p-value* less than 0.05 indicates statistical significance, and we applied a false discovery rate (FDR) for multiple comparisons to adjust statistical errors.

## Results

### Behavioural results

Reaction time (RT) distributions of internal/external trials in the introspection task are shown in Fig 2. There was a significant difference in RT between internal and external trials (*β* = -0.27,*p <*. *001*), indicating that participants were slower to respond to internal attention than external attention. This result is consistent with previous research, which used a similar approach (Hautekiet *et al*., 2023; Nishida *et al*., 2024), underscoring the validity of our experimental manipulation. However, there was a significant difference in RTs between repeated trials (in which the immediately previous trial was the same) and switched trials (in which the immediately previous trial was different) of internal attention (*β* = 0.038,*p <*. *001*), with slower RTs in switched trials than in repeated trials. RTs of switched trials were also significantly slower than repeated trials in external attention (*β* = 0.043,*p <*. *001*). This result is inconsistent with a previous study showing that switching costs were substantially asymmetric from external to internal attention (Verschooren *et al*., 2019).

**Figure 2.**
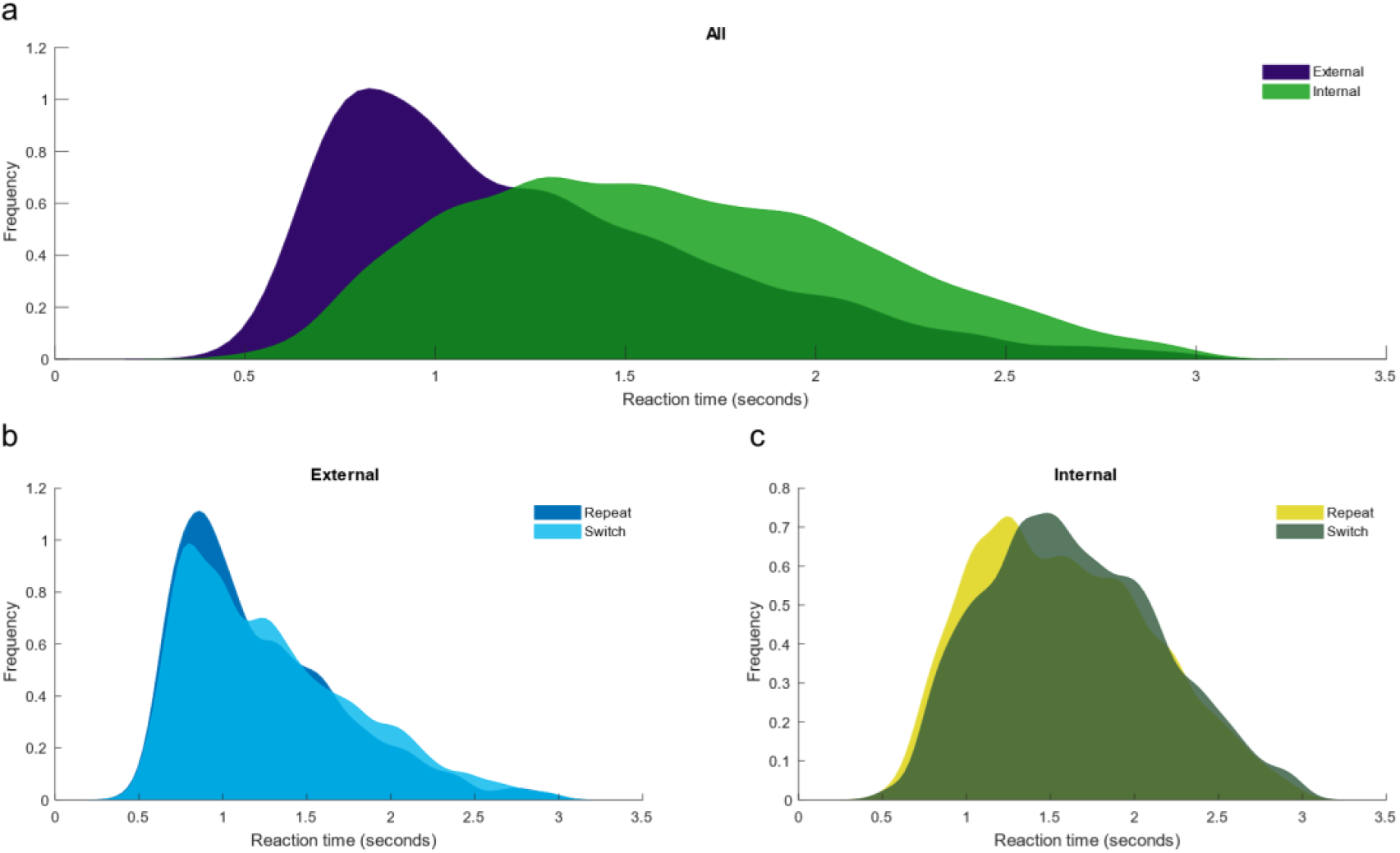
Differences in reaction time distributions among conditions. The figure was drawn using a kernel-fitted distribution (bandwidth: 0.08) for each direction (internal/external). **a. Reaction time distributions between attentional types**. Individual reaction times were pooled across participants in each condition. Those from external trials are coloured navy, and those from internal trials are coloured green. **b. Reaction time distributions between repeated and switched external trials**. Those from repeated trials are coloured blue, and those from switched trials are in cyan. **c. Reaction time distributions between repeated and switched internal trials**. Those from repeated trials are coloured yellow, and those from switched trials are in dark green.

### Dynamic dissimilarity of each cognitive process

After extracting the correlation matrix of each cognitive process (Fig 3a, for example), we obtained four DCIs corresponding to distance in attentional types (internal/external). All Wilcoxon test results were significant (*emotion*:*Z* = 3.72,*p*_*FDR*_ < 001,*r* = 0.88;*selective attention*:*Z* = 3.72,*p*_*FDR*_ < 001,*r* = 0.88;*self-referential thoughts*:*Z* = 3.72,*p*_*FDR*_ < 001,*r* = 0.88;*working memory*:*Z* = 3.33,*p*_*FDR*_ = 002,*r* = 0.79), which indicates that dynamics of these four cognitive processes differ significantly when attention is directed internally vs externally. We also confirmed these results using a permutation test that sorted the attentional type distance matrix and extracted DCI repeatedly (1000 times). We calculated the means of all participants in each permutation and compared them with the original means. Permutation test results were also significant for all cognitive processes (see Fig S3). There were also significant DCI differences between*emotion* and*working memory* (*Z* = 2.46,*p*_*FDR*_ = 0.028,*r* = 0.58). There were also differences between emotion vs. self-referential thoughts and selective attention vs. working memory, although those results did not survive after being corrected for multiple comparisons. All statistical values are shown in Table S2.

**Figure 3.**
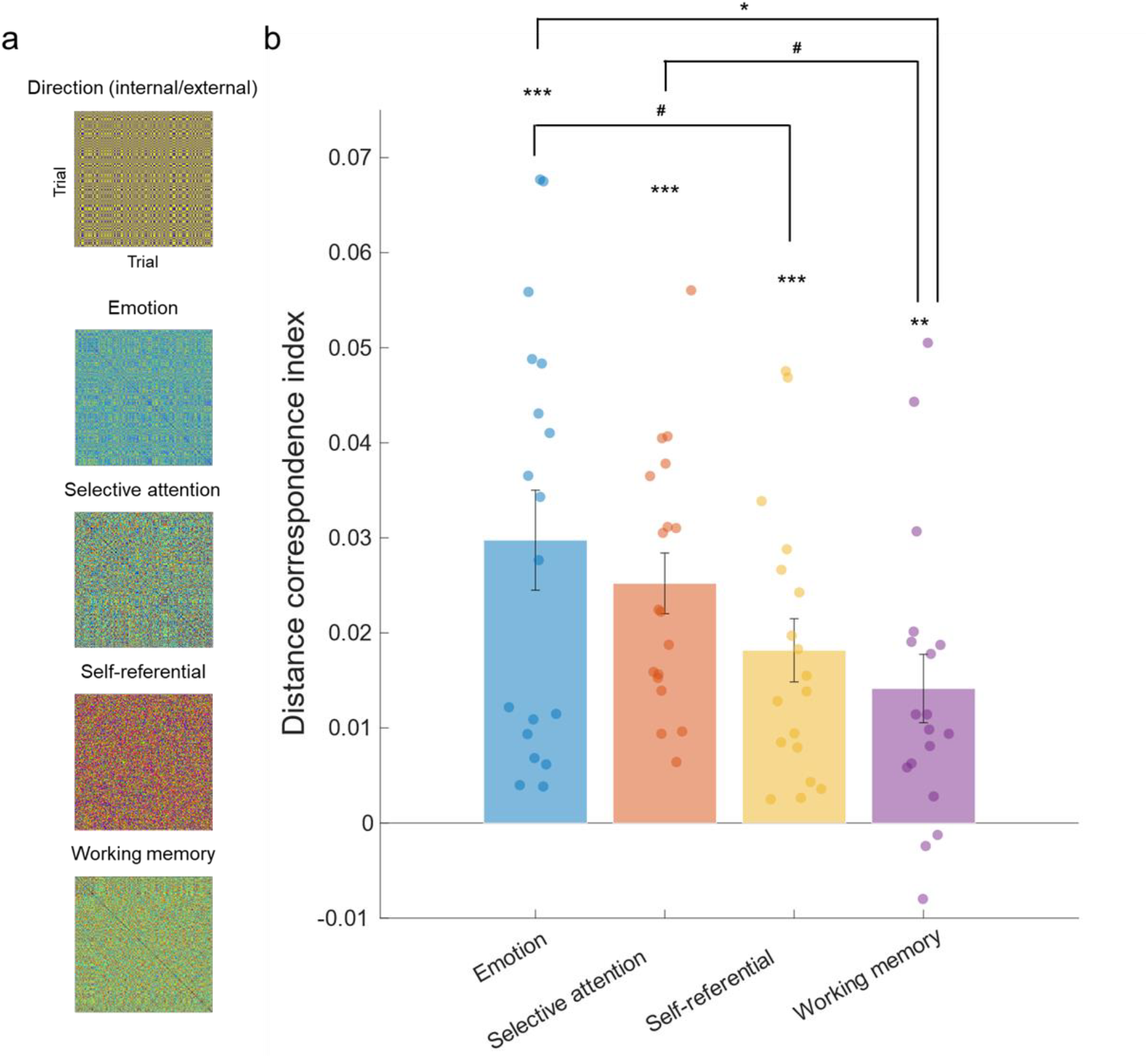
Comparison between dynamic dissimilarities of each cognitive process. **a.** Dynamic dissimilarity matrix from one example participant. The X-axis and Y-axis represent the trial index. GLM regression was done to compare the matrix of direction (attentional type) with each one of the cognitive processes, excluding no-interest regressors. **b**. Comparison of distance correspondence index (DCI) across cognitive processes. Error bars denote standard errors. #p < 0.05, uncorrected; *p < 0.05, **p < 0.01, ***p < 0.01, FDR-corrected for multiple comparisons over the number of statistical tests.

### Differentiation of each cognitive process was associated with clinical symptoms

There were significant negative associations between the DCI of selective attention and scores for depression and state anxiety (*β* = -0.002,*p*_*FDR*_ = 0.023;*β* = -0.002,*p*_*FDR*_ = 0.002), which indicates that greater similarity in selective attention between internal/external attention resulted in more severe symptoms. On the contrary, there was a significant positive association between the DCI of selective attention and the score of trait anxiety (*β* = 0.002,*p*_*FDR*_ = 0.002). Additionally, there was a positive association with ADHD in selective attention. There were positive associations with ADHD, state anxiety, and autism, and negative associations with gaming disorder and trait anxiety in self-referential. However, those did not survive multiple comparison corrections (see Fig. 4 and Table S3 for all statistical values).

**Figure 4.**
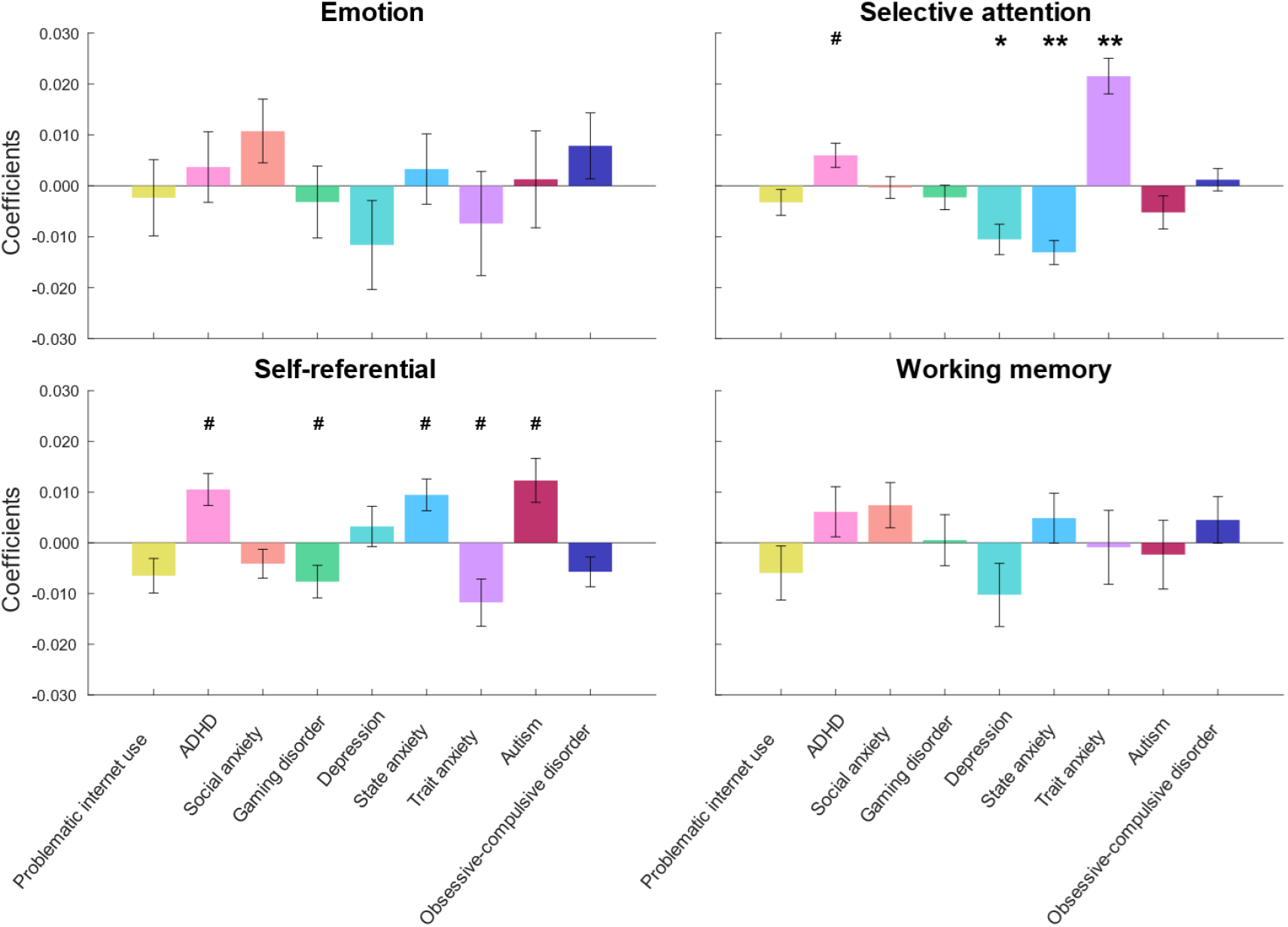
Associations between DCI in each cognitive process and psychiatric symptoms. Error bars denote standard errors. #p < 0.05, uncorrected; *p < 0.05, **p < 0.01, FDR-corrected for multiple comparisons over the number of psychiatric independent variables investigated.

## Discussion

In this study, we used dynamic similarity analysis of fMRI data captured during an introspection task to reveal differences in cognitive processes associated with internal/external attention. Specifically, we examined whether temporal dynamics of four global cognitive functions (emotion processing, selective attention, self-referential thoughts, and working memory) significantly differed between internal/external attention. Our results indicate that validating the nature of the task manipulation, emotion function appears to differ more strongly than other cognitive processes, such as working memory. In addition, we tested whether these cognitive differences between internal/external attention were associated with several psychiatric disorder scores. We show preliminary results that the dissimilarity of selective attention was negatively associated with depression and state anxiety, and positively associated with trait anxiety scores.

All extracted cognitive dynamics differed significantly between internal and external attention states, with particularly robust differences between internal and external attention states in emotional processing. This is a reasonable result given the task’s characteristic focus on emotion versus letter counts. This finding suggests that the degree of each cognitive dimension dynamic in our brain differs depending on the object/content being processed.

However, the present results may simply depend on our task paradigm. For example, some studies have conducted similar tasks related to internal and external attention. Still, participants were asked to perform a (working) memory search instead of focusing on emotions during internal attention (Verschooren *et al*., 2019). In such tasks, degrees of difference in each cognitive process might differ from our results. Indeed, such differences in processes across task types may be reflected in the fact that results of our reaction time analysis are not consistent with previous findings using different tasks. Therefore, future work should consider the generalisability of results across task types.

Regression analyses related to psychiatric disorders indicate there is a clear relationship between the similarity of selective attention, but not in emotion and several psychiatric severities. Our results show that only depression and anxiety are related to selective attention dynamics in the brain. These findings provide mechanistic evidence for a greater effect on depression and anxiety compared to other disorders by cognitive interventions such as Attention Training Technique (ATT), to improve attention control related to psychiatric symptoms, e.g., rumination (Knowles *et al*., 2016). Moreover, interesting opposing effects between depression/state anxiety and trait anxiety were found in the association with the similarity of selective attention. While the result that excess similarity of selective attention dynamics between internal and external attention is associated with state anxiety follows our a priori hypothesis, the relationship is inverse with respect to trait anxiety. Considering previous evidence for differences between anxiety types in attentional functions (Pacheco-Unguetti *et al*., 2010) and brain network levels (Saviola *et al*., 2020), this contradictory result may reflect an overreaction of attentional control in people with high trait anxiety. In light of this issue, when considering characteristics of trait anxiety, it might be better to adopt interventions that balance the direction of attention for high-trait anxious patients.

The study has several other limitations. First, the sample size is relatively low. While we acknowledge the limited sample size (N=18), repeated measures within a participant and our advanced data analysis based on trial-wise level allowed more fine-grained and informative insights for cognitive processes than traditional fMRI studies, making this small sample size sufficient to detect meaningful patterns (Nee, 2018). Second, cognitive functions used in C-DSA were selected a priori based on the hypothesis, but it is possible to apply data-driven sampling (Koyama *et al*., 2023) or task-based activation maps (Kragel *et al*., 2023). Although we had explicit hypotheses based on previous theories, it is possible that using different approaches might lead to different results. Third, although we focused on symptoms of each psychiatric disorder independently, a transdiagnostic perspective may lead to different relationships (Gillan *et al*., 2016; Seow & Gillan, 2020). In the future, one could examine the relationship with such transdiagnostic factors using factor analysis results from large-scale data (Wise *et al*., 2023). Fourth, participants were healthy, and we did not include patients undergoing psychiatric treatment. Generalisability of these results in clinical populations should be investigated.

### Methodological implications: The temporal window of cognitive processes in the brain

C-DSA allows us to quantify temporal dynamics of cognitive processes in complex functions. Though previous research has already suggested dynamic representational similarity analysis to assess the temporal neural representation in the brain (de Vries & Wurm, 2023), it cannot be applied to clarify what and how cognitive properties are associated with complex naturalistic processes.

Moreover, C-DSA can be expanded to both naturalistic and (pseudo)controlled dynamic processes, which are considered constructed from complex systems. Though we used the C-DSA in a pseudo-controlled experiment without substantial restrictions on reaction time, the technique could be applied to experiments using more naturalistic stimuli and settings, e.g., while watching movies, social communication, etc.

Moreover, as shown here, C-DSA enables us to clarify the association between complex “dynamic” processes and several factors, such as ageing and psychiatric disorders. Though some previous research attempted to clarify how neural differentiation is associated with such problems (Katsumi *et al*., 2021; Tamman *et al*., 2024), those results only provided knowledge related to concrete structures and functions of the brain that do not help clinical practice directly. C-DSA can provide more clinically oriented evidence and can suggest which cognitive processes are involved in modulating psychiatric symptoms with regard to complex cognitions/behaviours. Taken together, C-DSA could open a new horizon to explore dynamic cognitive mechanisms underlying complex functions across a wide range of levels. In this sense, it could contribute to reorganising such inconsistently defined concepts as internal/external attention from the theoretical and philosophical perspective of cognitive ontology (Poldrack & Yarkoni, 2016).

## Conclusion

In summary, by comparing time series dynamics of each cognitive process in internal/external attention tasks using C-DSA, we showed that each cognitive function may have different dynamics among types of attention. More notably, estimated similarities between time series may contain information on the temporal characteristics of emotional and cognitive functions related to psychiatric symptom severity. These results highlight the potential of C-DSA not only to help reveal complex cognitive dynamics, such as internal/external attention, but also to provide insights into cognitive processes underlying psychiatric disorders (Millan *et al*., 2012).

## Data availability

Relevant data, scripts, and materials are available in OSF (https://osf.io/cqknz/).

## Declaration of AI use

We used AI-assisted technologies while drafting the manuscript to improve the readability and language of the work.

## Author contributions

TO designed the study with inputs from AC and MM, acquired data, analysed data, and prepared the original draft. YK supervised the analytical process. YK, AS, MM, NK, and AC reviewed and edited the manuscript. All authors approved this submission and agreed to take responsibility for the manuscript.

## Acknowledgements

We thank Kaori Nakamura for recruiting and collecting data and Naoyuki Okamoto for advice about methodology.

