## Supplementary materials for "Dynamic cognitive differences between internal and external attention are associated with depressive and anxiety disorders"

### Supplementary methods

#### The details of the introspection task

Instructions given participants were as follows:

"You will now see several questions and words on the screen in front of you. For the question 'How much do you feel?' please indicate the degree to which you feel the emotion shown, and for the question 'How many letters are there?' please indicate the number of letters in the word displayed. Please be careful not to press the incorrect answer. Please answer honestly, focusing on the number of letters and the feeling of the word at that moment, rather than choosing the same response all the time."

A list of stimuli representing mood, extracted from EACL (Oda et al., 2015) is shown in Table S1. To match the number of negative-positive words, we took one or more from each factor of the negative questions and confirmed that the average number of letters matched. An example of actual Japanese word stimuli is shown in Fig S1.

**Table S1 The list of stimuli used in the task**

| Positive | Negative |
|---|---|
| 不安な - Anxious | 楽しい - Fun |
| びくびくしている - Nervous | さわやかな - Refreshing |
| いらいらしている - Irritated | 快い - Pleasant |
| 気が立っている - Agitated | 満ち足りた - Content |
| 落ち込んでいる - Depressed | 気力に満ちた - Enthusiastic |
| ふさいでいる - Gloomy | 活動的な - Active |
| 悲しい - Sad | 機敏な - Alert |
| 不快な - Uncomfortable | エネルギッシュな - Energetic |
| 嫌な - Unpleasant | ゆったりした - Relaxed |
| 気分ののらない - Unmotivated | 落ち着いた - Calm |
| 気が休まらない - Restless | くつろいだ - Comfortable |

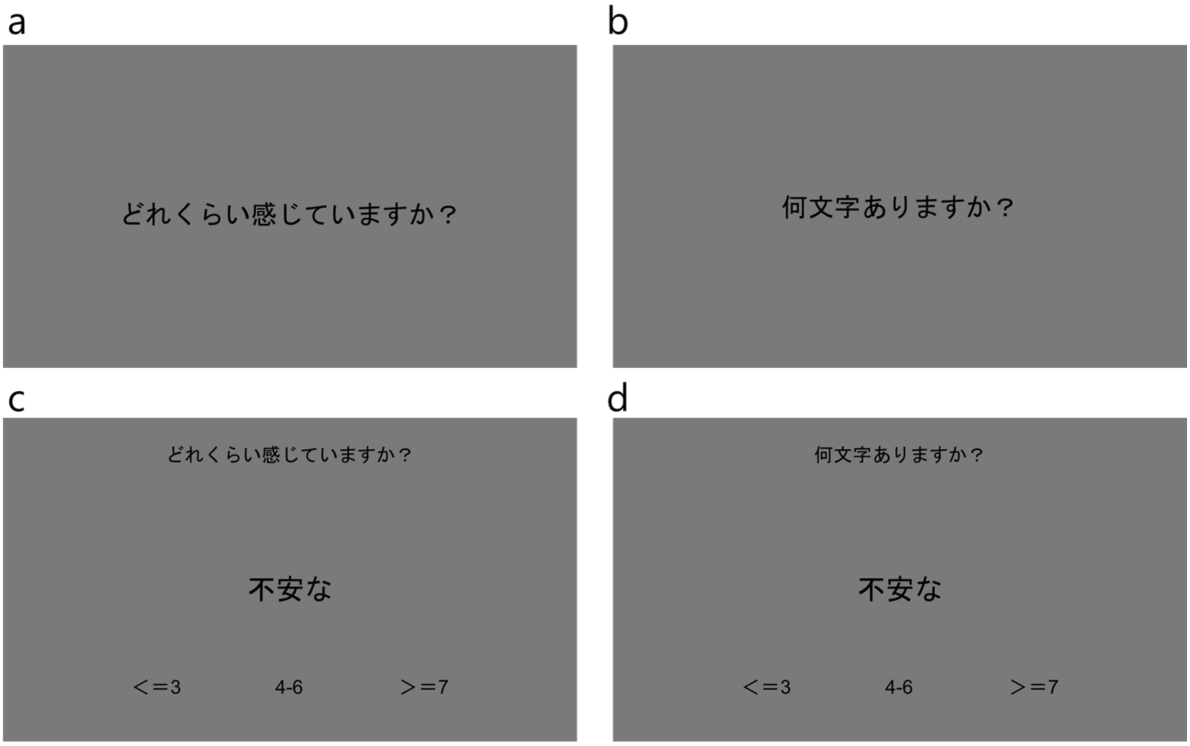


**Figure S1 Examples of instruction stimuli from each internal/external attentional trial**

*(a) The first instruction for internal attention ("How much do you feel?") (b) The first instruction for external attention ("How many letters?") (c)(d) An example of an emotional word prompt ("anxious").*

#### Details of questionnaires used for screening and assessment

Major depressive disorder was assessed using the Beck Depression Inventory II (BDI-II), which demonstrated an internal consistency of 0.91 in its original version ((Dozois *et al.*, 1998)). Construct validity was confirmed using the original BDI ($r$ = 0.93) ((Dozois *et al.*, 1998)). Its reliability and validity have been verified in Japanese populations (Cronbach's α = 0.87) (Kojima *et al.*, 2002). General state-trait anxiety was evaluated with the State-Trait Anxiety Inventory (STAI) (Spielberger, 1983), which reported a Cronbach's alpha ranging from 0.86 to 0.95 in the original study. The Japanese version also showed confirmed reliability and validity (Cronbach's α = 0.92) (Iwata *et al.*, 1998). Social anxiety was measured using the Liebowitz Social Anxiety Scale (LSAS-fear/avoid) (Baker *et al.*, 2002), with a Cronbach's alpha of 0.95 in the original version. Construct validity was established with Safren's four-factor model in the original study (Baker *et al.*, 2002) and validated against the Social Avoidance and Distress Scale and professional diagnosis in the Japanese version (Asakura *et al.*, 2002). Obsessive-compulsive disorder (OCD) was assessed using the Obsessive-Compulsive Inventory (OCI), which has a Cronbach's alpha between 0.86 and 0.95 in the original version (Foa *et al.*, 1998) and 0.96 in the Japanese version (Ishikawa *et al.*, 2014). Construct validity was confirmed with the Yale-Brown Obsessive Compulsive Scale, Compulsive Activity Checklist, Maudsley Obsessive-Compulsive Inventory (r = 0.74), STAI (r = 0.41), and Center for Epidemiologic Studies Depression Scale (r = 0.48) in the Japanese version (Ishikawa *et al.*, 2014). Internet-related problems were evaluated using the Compulsive Internet Use Scale (CIUS), with a Cronbach's alpha of 0.89 in the original version (Meerkerk et al., 2009) and over 0.9 in the Japanese version (Yong et al., 2017). Construct validity was demonstrated through strong positive correlations with the Online Cognition Scale (r = 0.70, p < 0.001) and the amount of time spent online (r = 0.33, p < 0.001) in the original version, and POSI, MR, DSR, NO, K6, and UCLA Loneliness in the Japanese version. Autism Spectrum Disorder (ASD) was measured with the Autism-Spectrum Quotient (AQ), showing a Cronbach's alpha between 0.63 and 0.77 in the original version (Baron-Cohen *et al.*, 2001) and an internal consistency of 0.81 in the Japanese version (Wakabayashi *et al.*, 2006). Construct validity was confirmed by professional diagnosis based on DSM-IV (r = 0.58) and exposure to social adaptation problems (r = 0.92) in the Japanese version (Wakabayashi *et al.*, 2006). Adult attention-deficit/hyperactivity disorder (ADHD) was assessed using the Adult ADHD Self-Report Scale (ASRS) (Kessler *et al.*, 2005), which had a Cronbach's alpha between 0.88 and 0.89 in the original version (Adler *et al.*, 2006), and 0.83 in the Japanese version (Takeda *et al.*, 2017). Construct validity was verified against the Japanese version of the Conners' Adult ADHD Rating Scales-Self Report and BDI-II (Takeda *et al.*, 2017). The Internet Gaming Disorder Scale (IGDS) includes questions corresponding to each of the nine gaming disorder symptoms defined in DSM-5 (American Psychiatric Association, 2013). Using a binary response format, it assesses the presence of each symptom during the preceding year (Lemmens *et al.*, 2015). A full-syndrome diagnosis requires at least five symptoms, while three symptoms indicate a sub-threshold gaming disorder (Lemmens *et al.*, 2015). Reliability and validity have been confirmed (Lemmens *et al.*, 2015), and the questionnaire has proven effective for screening gaming disorder (King *et al.*, 2020). The Japanese version has also demonstrated reliability and validity (Sumi *et al.*, 2018).

### Supplementary Results

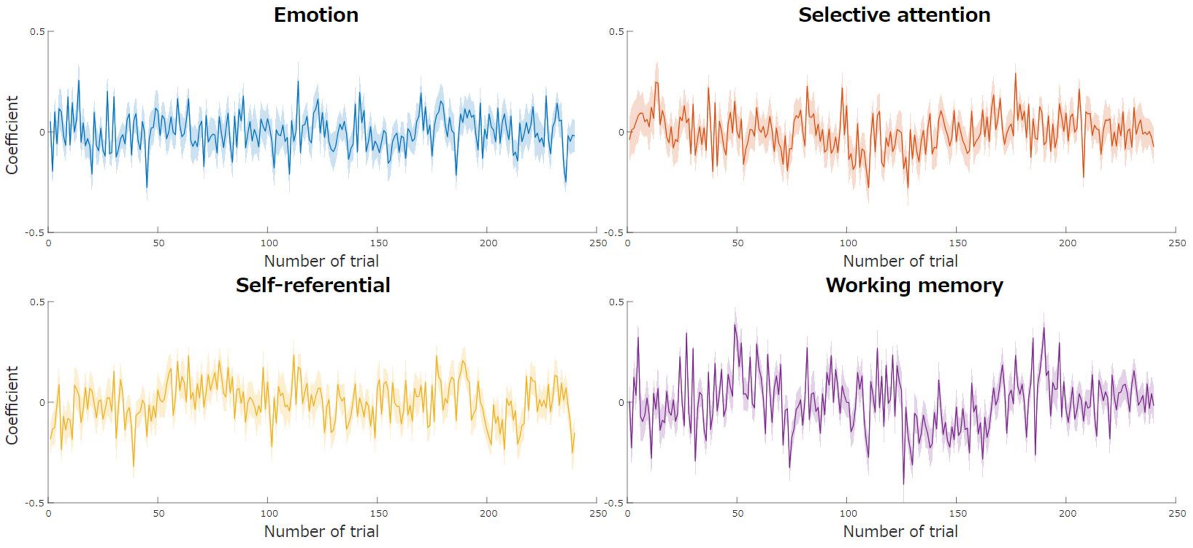


**Figure S2. The time series of each cognitive process through sessions**

*Coefficients are averaged across all sessions of all participants. The shaded area indicates the standard error. Note that averaged coefficients across participants, which are shown here, may have reflected only noise components of the experiment and may not have included specific internal/external attentional processes.*

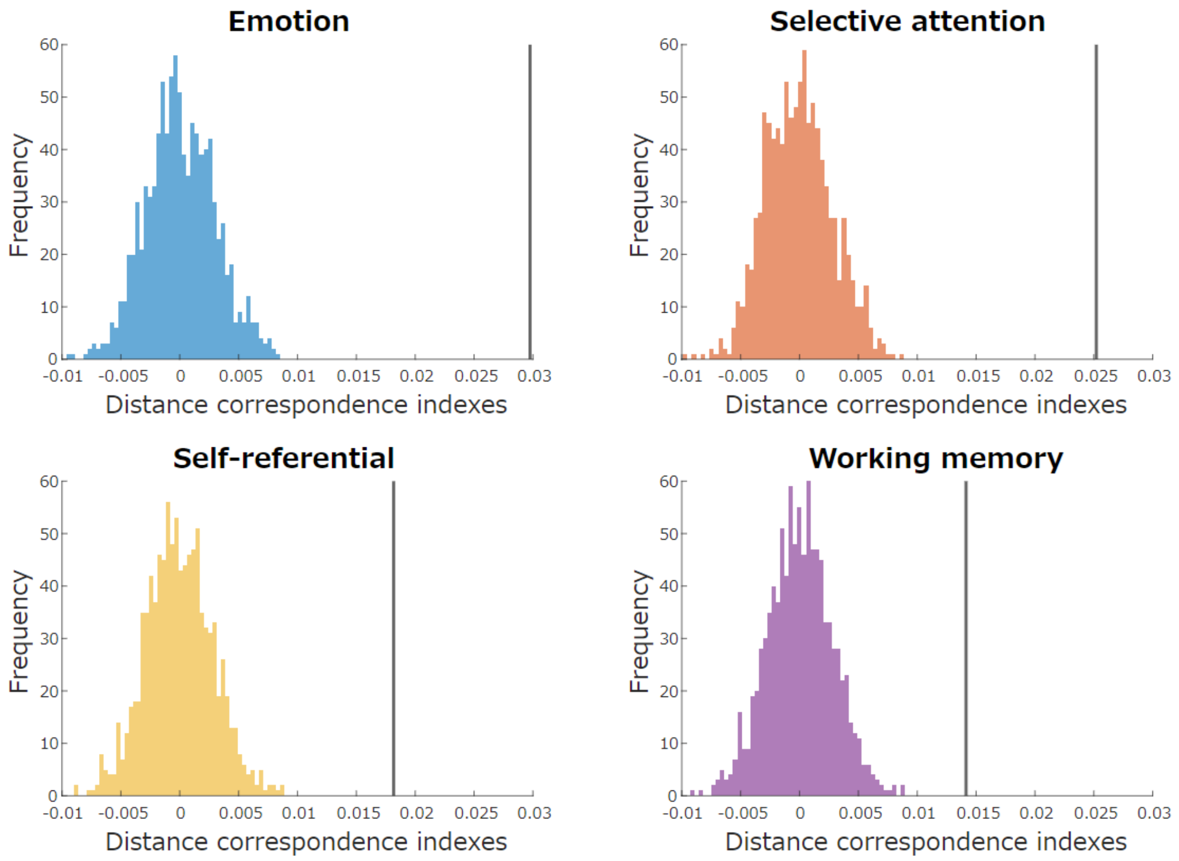


**Figure S3. Distance correspondence indices of cognitive processes**

*Vertical solid lines represent the average value of the original distance correspondence index from our sample (N=18). All average values differed significantly from the distribution of random permutations (1000 times).*

| | $Z$ | $p_{FDR}$ | $r$ |
|---|---|---|---|
| Emotion | 3.724 | < 0.001 | 0.878 |
| Selective attention | 3.724 | < 0.001 | 0.878 |
| Self-referential | 3.724 | < 0.001 | 0.878 |
| Working memory | 3.332 | 0.002 | 0.785 |
| Emotion vs Selective attention | 0.152 | 0.879 | 0.036 |
| Emotion vs Self-referential | 1.851 | 0.092 | 0.436 |
| Emotion vs Working memory | 2.461 | 0.028 | 0.580 |
| Selective attention vs Self-referential | 1.502 | 0.166 | 0.354 |
| Selective attention vs Working memory | 2.112 | 0.058 | 0.498 |
| Self-referential vs Working memory | 0.936 | 0.388 | 0.221 |

**Table S2. Sign-ranked test for each cognitive process**

| | **Emotion** | | | **Selective attention** | | | **Self-referential** | | | **Working memory** | | |
|---|---|---|---|---|---|---|---|---|---|---|---|---|
| | Beta | SE | $p_{FDR}$ | Beta | SE | $p_{FDR}$ | Beta | SE | $p_{FDR}$ | Beta | SE | $p_{FDR}$ |
| Problematic internet use | -0.00030 | 0.00097 | 0.858 | -0.00042 | 0.00033 | 0.359 | -0.00084 | 0.00044 | 0.119 | -0.00077 | 0.00069 | 0.534 |
| ADHD | 0.00035 | 0.00065 | 0.853 | 0.00056 | 0.00022 | 0.079 | 0.00099 | 0.00030 | 0.065 | 0.00058 | 0.00046 | 0.534 |
| Social anxiety disorder | 0.00043 | 0.00025 | 0.778 | -0.00001 | 0.00009 | 0.875 | -0.00017 | 0.00011 | 0.209 | 0.00030 | 0.00018 | 0.534 |
| Gaming disorder | -0.00205 | 0.00455 | 0.853 | -0.00146 | 0.00155 | 0.481 | -0.00493 | 0.00207 | 0.080 | 0.00033 | 0.00324 | 0.920 |
| Major depressive disorder | -0.00164 | 0.00123 | 0.778 | -0.00149 | 0.00042 | 0.023 | 0.00046 | 0.00056 | 0.439 | -0.00145 | 0.00088 | 0.534 |
| State-anxiety | 0.00037 | 0.00078 | 0.853 | -0.00147 | 0.00027 | 0.002 | 0.00106 | 0.00035 | 0.065 | 0.00055 | 0.00055 | 0.534 |
| Trait-anxiety | -0.00077 | 0.00106 | 0.853 | 0.00223 | 0.00036 | 0.002 | -0.00122 | 0.00048 | 0.079 | -0.00009 | 0.00075 | 0.920 |
| Autism | 0.00021 | 0.00152 | 0.896 | -0.00084 | 0.00052 | 0.263 | 0.00197 | 0.00069 | 0.065 | -0.00037 | 0.00108 | 0.920 |
| Obsessive-compulsive disorder | 0.00043 | 0.00035 | 0.778 | 0.00007 | 0.00012 | 0.677 | -0.00031 | 0.00016 | 0.119 | 0.00024 | 0.00025 | 0.534 |

**Table S3. Linear regression of each DCI and psychiatric scores**
